# Retention of ES cell-derived 129S genome drives NLRP1 hypersensitivity and transcriptional deregulation in *Nlrp3^−/−^* mice

**DOI:** 10.1101/2024.01.03.573991

**Authors:** Felix D. Weiss, Yubell Alvarez, Anshupa Sahu, Farhad Shakeri, Hye Eun Lee, Anne-Kathrin Gellner, Andreas Buness, Eicke Latz, Felix Meissner

## Abstract

Immune response genes are highly polymorphic in humans and mice, with heterogeneity amongst loci driving strain-specific host defense responses. The inadvertent retention of polymorphic loci can introduce confounding phenotypes, leading to erroneous conclusions, and impeding scientific advancement. In this study, we employ a combination of RNAseq and variant calling analyses and identify a substantial region of 129S genome, including the highly polymorphic *Nlrp1* locus proximal to *Nlrp3*, in one of the most commonly used mouse models of NLRP3 deficiency. We show that increased expression of 129S NLRP1b sensitizes *Nlrp3*^−/−^ macrophages to NLRP1 inflammasome activation. Furthermore, the presence of 129S genome leads to altered gene and protein regulation across multiple cell-types, including of the key tissue-resident macrophage marker, TIM4. To address the challenge of resolving NLRP3-dependent phenotypes, we introduce and validate a conditional *Nlrp3* allele, enabling precise temporal and cell-type-specific control over *Nlrp3* deletion. Our study establishes a generic framework to identify functionally relevant SNPs and assess genomic contamination in transgenic mice. This allows for unambiguous attribution of phenotypes to the target gene and advances the precision and reliability of research in the field of host defense responses.

## Introduction

Evolutionary pressure results in the emergence of gene paralogs and polymorphisms that shape protein function and regulation. Due to strong selection pressure, immune genes are disproportionately hyperpolymorphic across inbred mouse strains^1^, resulting in strain-specific host defense mechanisms, subsequently maintained through generations of inbreeding.

The retention of embryonic stem cell (ESC)-derived genetic material in transgenic mice, especially of polymorphic immune loci, can lead to confounding phenotypes independent of the target gene^2–5^, as well as the identification of novel immune defense mechanisms^6^.

Inflammasomes are key innate immune signaling hubs that when activated induce a lytic form of cell death known as pyroptosis, and the release of the inflammatory cytokines IL-1β and IL-18. While the inflammasome protein NLRP3 is activated by danger signals including viral infection, potassium efflux and excessive extracellular ATP, the murine NLRP1 inflammasome can be activated by Anthrax lethal toxin, *Toxoplasma gondii* (*T. gondii*) infection, and inhibition of dipeptidyl proteases (DPP) 8/9. Whether an endogenous murine NLRP1 activator exists remains unknown.

While, NLRP3 is largely conserved across inbred laboratory mouse strains, the neighboring *Nlrp1* locus is highly variable^7,8^. There are five *Nlrp1* genes of which, *Nlrp1a*, *Nlrp1b* and *Nlrp1c* are present in C57B6 strains, and *Nlrp1b* and *Nlrp1c* in 129S strains^8^. Furthermore, five distinct *Nlrp1b* alleles, distributed across 14 laboratory mouse strains, display differential sensitivities to a diverse range of stimuli^7^, and may be subject to alternative methods of regulation. Studying strain-specific sequence differences in *Nlrp1b* that may contribute to its alternative sensitivities or regulation has relied largely on the generation of overexpressing cell lines, or whole animals where genetic differences are not limited just to *Nlrp1b*, and can contribute to observed phenotypes. A maximally genetically homogeneous model system, where functionally relevant cells such as macrophages, express different *Nlrp1b* alleles, controlled by their endogenous regulatory elements, would provide a valuable tool to further our understanding of NLRP1 biology.

NLRP3 has been implicated in a large number of diseases. Murine models of NLRP3 deficiency have shown NLRP3’s causal contribution to the pathogenesis of gout^9^ and atherosclerosis^10^, through activation by uric acid crystals and cholesterol crystals respectively, however its mechanistic contribution to diseases including multiple sclerosis^11,12^, Alzheimer’s disease^13,14^, and diet induced inflammation^15^ remains to be clarified. Previous research has shown beneficial effects on disease phenotypes in *Nlrp3^−/−^*mice even in the absence of clear evidence for NLRP3 inflammasome activation in vivo^12–18^, and with limited pharmacological validation. This suggests a potential inflammasome independent role for NLRP3, or an animal model effect independent of NLRP3 deficiency.

Our study identifies a substantial region of 129S ESC-derived genome in a frequently used model of *Nlrp3* deficiency (Nlrp3KO^129ES^, Ref. ^19^). The ∼40Mb region on chromosome 11 contains several hundred genes critical for cell function and identification, and key immune genes, including the *Nlrp1* locus. We leverage this unexpected occurrence to identify strain-specific differences that influence the NLRP1 inflammasome response and its post-translational regulation, in a largely genetically homogenous model. Differences in protein coding sequences identified in this study impact innate immune responses independently of the loss of NLRP3 expression, and need to be considered when ascribing mechanistic phenotypes in Nlrp3KO^129ES^ mice. Validation of a novel inducible *Nlrp3* allele allowing for temporal and cell-type specific control of *Nlrp3* deletion will provide greater clarity on the mechanistic contribution of NLRP3 to disease pathology, in the absence of confounding effects. Finally, our analytical strategy to identify coding variants in relevant expressed genes is applicable to historic and newly generated datasets, enabling a straight-forward analysis of coding variants and genetic heterogeneity in transgenic mice currently considered congenic.

### Nlrp3KO^129ES^ mouse macrophages display increased sensitivity to NLRP1 activation by Talabostat

DPP8/9 inhibition by Talabostat activates the NLRP1 inflammasome (**Fig 1A**, Ref. ^20–23)^. We compared the dose-dependent activation of NLRP1 by Talabostat in bone marrow derived macrophages (BMDMs), generated from C57B6/J and Nlrp3KO^129ES^ mice, backcrossed >10 generations to C57B6/J and defined by Charles River as congenic.

**Figure 1.**
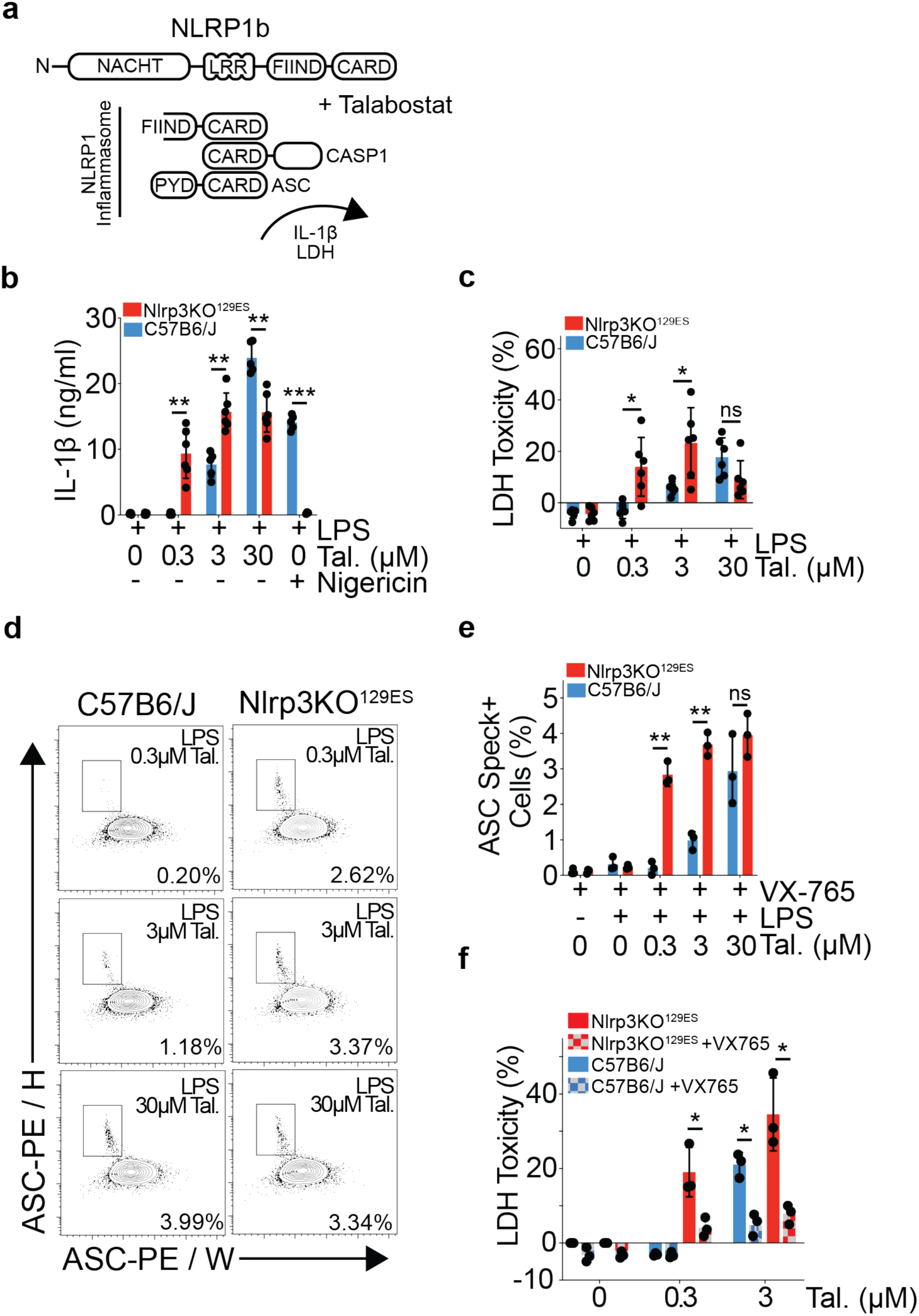
Nlrp3KO^129ES^ mouse macrophages are hypersensitive to NLRP1 inflammasome activation by Talabostat. **a.** Schematic showing murine NLRP1 inflammasome formation in response to Talabostat treatment. **b.** IL-1β release from C57B6/J and Nlrp3KO^129ES^ mouse BMDMs stimulated with LPS (10 ng/ml) and Talabostat (0.3 μM, 3 μM or 30 μM) for 24 hours, or primed with LPS (10 ng/ml) for 3 hours and then stimulated with Nigericin (8 μM) for 90 minutes (C57B6/J, n = 5, Nlrp3KO^129ES^, n = 6.) **c.** LDH release, relative to untreated total lysis controls, measured from C57B6/J and Nlrp3KO^129ES^ mouse BMDMs stimulated with LPS (10 ng/ml) and Talabostat (0.3 μM, 3 μM or 30 μM) for 24 hours (C57B6/J, n = 6, Nlrp3KO^129ES^, n = 6). **d.** Representative flow cytometry plots showing identification of ASC speck positive cells in C57B6/J and Nlrp3KO^129ES^ mouse BMDMs, pre-gated on live single cells, stimulated with LPS (10 ng/ml) and Talabostat (0.3 μM, 3 μM or 30 μM) for 16 hours, in the presence of VX-765 (50 μM). Percentages represent % of ASC Speck+ cells in the representative plot. **e.** Bar plot showing % of ASC speck positive BMDMs from C57B6/J and Nlrp3KO^129ES^ mice stimulated with LPS (10ng/ml) and Talabostat (0.3 μM, 3 μM or 30 μM) for 16 hours, in the presence of VX-765 (50 μM, C57B6/J, n = 3, Nlrp3KO^129ES^, n = 3). **f.** LDH release, relative to total lysis controls, measured from C57B6/J and Nlrp3KO^129ES^ mouse BMDMs stimulated with Talabostat (0.3 μM or 3 μM) in the presence or absence of VX-765 (50 μM) for 24 hours (C57B6/J, n = 3, Nlrp3KO^129ES^, n = 3). **b - f.** All *P* values were calculated using multiple unpaired parametric t-tests. FDR (*q*) was calculated using Benjamini Hochberg correction. * = *q* < 0.05, ** = *q* < 0.01, *** = *q* < 0.001, **** = *q* < 0.0001, ns = not significant (*q* > 0.05). Error bars represent standard deviation.

Nlrp3KO^129ES^ versus C57B6/J BMDMs showed a significant increase in IL-1β secretion at low doses of Talabostat stimulation (**Fig 1B**). Consistent with a lower threshold of NLRP1 inflammasome formation, the supernatant LDH levels – indicative of cell membrane rupture by pyroptotic cell death – were also significantly higher in Nlrp3KO^129ES^ BMDMs (**Fig 1C**). These data show that Nlrp3KO^129ES^ BMDMs are hypersensitive to DPP8/9 inhibition induced IL-1β release and pyroptosis compared to C57B6/J controls.

Activation of the NLRP1 inflammasome leads to ASC speck formation, where Caspase-1 is recruited and activated. The frequency of ASC speck positive cells was significantly higher at lower doses of Talabostat treatment in Nlrp3KO^129ES^ BMDMs (**Fig 1D, E**). These results mirror those observed by IL-1β and LDH release, and show a lower threshold for inflammasome assembly upon NLRP1 activation in Nlrp3KO^129ES^ versus C57B6/J BMDMs. Caspase-1 inhibition by VX-765 significantly rescued cell death, as measured by LDH, in both genotypes (**Fig 1F**), showing that increased IL-1β and LDH release is due to the formation of a functional inflammasome.

### NLRP1b protein, but not transcript, is upregulated in Nlrp3KO^129ES^ BMDMs

In order to determine whether the increased sensitivity to NLRP1 activation in Nlrp3KO^129ES^ BMDMs was related to NLRP1b protein expression levels, we measured protein abundance by LC-MS/MS. As expected, NLRP3 was not detected in Nlrp3KO^129ES^ BMDMs (**Fig 2A; Supplemental Table 1**). Conversely, the NLRP1 inflammasome forming protein, NLRP1b, was not detected in Nlrp3WT^129ES^ BMDMs, but was robustly expressed at baseline, and upregulated in response to 24 hours of LPS stimulation, in Nlrp3KO^129ES^ BMDMs (**Fig 2A**).

**Figure 2.**
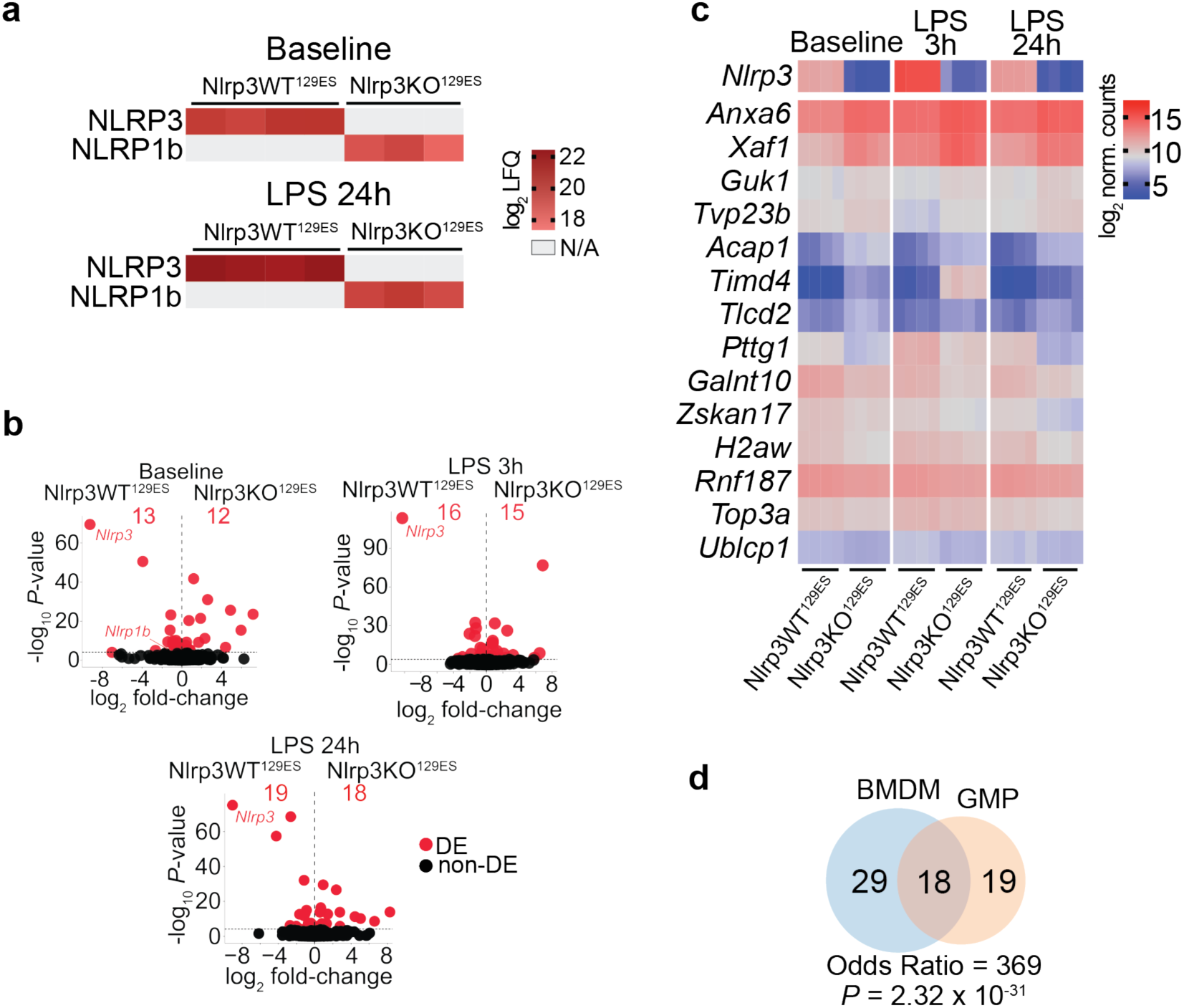
Nlrp3KO^129ES^ macrophages have increased NLRP1b protein expression, with limited global gene expression changes. **a.** Heatmap of log_2_ LFQ values of NLRP3 and NLRP1b proteins from LC/MS-MS of Nlrp3WT^129ES^ and Nlrp3KO^129ES^ BMDMs at baseline and after 24 hours of LPS (10 ng/ml) stimulation. Grey spaces indicate samples where NLRP3 or NLRP1b were not detected. **b.** Volcano plots of gene expression fold-change vs. *P*-value in RNA-seq of Nlrp3WT^129ES^ and Nlrp3KO^129ES^ BMDMs. Total number of differentially expressed (DE, adj. *P* < 0.05, Benjamini-Hochberg adjusted) genes, and individual DE genes, are shown in red. Left: Comparison at baseline. Right: Comparison after 3 hours of LPS (10 ng/ml) stimulation. Bottom: Comparison after 24 hours of LPS (10 ng/ml) stimulation. **c.** Heatmap of log2 normalised counts of genes commonly differentially expressed (adj *P* < 0.05, Benjamini-Hochberg adjusted) in all three comparisons above. **d.** Venn diagram showing enrichment of shared DE genes (adj. *P* < 0.05) between Nlrp3WT^129ES^ and Nlrp3KO^129ES^ in BMDMs and GMPs. One-sided Fisher’s exact test was applied for the odds ratio and *P* value.

In order to understand if NLRP1b protein upregulation in Nlrp3KO^129ES^ BMDMs was transcriptionally dependent, we performed RNA sequencing (RNAseq) on polyA-RNA from BMDMs from Nlrp3KO^129ES^ and Nlrp3WT^129ES^ littermate controls. Gene expression analysis showed minor changes between Nlrp3KO^129ES^ and Nlrp3WT^129ES^ BMDMs at baseline and after LPS stimulation (**Fig 2B, Supplemental Fig 1; Supplemental Table 2**). Unexpectedly, *Nlrp1b* was slightly upregulated in Nlrp3WT^129ES^ BMDMs at baseline, however this differential expression was not observed following LPS stimulation (**Fig 2B**). The remaining deregulated genes showed no statistically significant shared functional enrichment as determined by Gene Ontology analysis of biological process or molecular function (adj. *P* < 0.05), and included genes related to annexins (*Anxa6*), apoptosis regulation (*Xaf1*), cell cycle (*Pttg1*) and histones (*H2aw*) among others (**Fig 2C**).

These data show that Nlrp3KO^129ES^ BMDMs upregulate NLRP1b protein expression, independently of an increase in *Nlrp1b* transcript. Furthermore, Nlrp3KO^129ES^ BMDMs show a limited number of differentially expressed genes with no enriched pathways or functions.

### Gene expression changes are shared between BMDMs and granulocyte monocyte progenitors in Nlrp3KO^129ES^ mice

Next, we asked whether gene expression changes also occur in non-differentiated myeloid cells. BMDMs are a model of monocyte-derived macrophages that arise from precursors in the bone marrow, including granulocyte-monocyte progenitors (GMPs). We performed gene expression analysis on Nlrp3KO^129ES^ and Nlrp3WT^129ES^ litter-mate control GMPs, isolated by FACS from murine bone marrow (**Supplemental Fig 2A**). Similar to Nlrp3KO^129ES^ BMDMs, there was a limited number of significantly differentially expressed genes (**Supplemental Fig 2B, C; Supplemental Table 3**).

Strikingly, despite the limited number of significantly differentially expressed genes, there was highly significant overlap between the two cell-types (48% of DE genes in GMPs, 38% of DE genes in BMDMs, **Fig 2D**). Commonly deregulated genes included but were not limited to *Xaf1*, *Anxa6, Pttg1*, *H2aw* and *Nlrp1b* (**Supplemental Fig 2D**).

### Genes differentially expressed in Nlrp3KO^129ES^ BMDMs and GMPs, including *Nlrp1b*, are predominantly located on chromosome 11, and are of 129S origin

Analysis of chromosomal location of deregulated genes in Nlrp3KO^129ES^ BMDMs and GMPs, revealed that deregulated genes were highly enriched on Chromosome 11, with 45 out of 47 (95.7%) significantly deregulated genes in BMDMs and 35 out of 37 (94.6%) in GMPs (**Fig 3A**). Strikingly, deregulated genes in BMDMs on chromosome 11 were located within a confined region +/− 25Mb proximal to *Nlrp3*. The Nlrp3KO^129ES^ mice used in this study were originally generated in 129SvEvBRD Lex1 ESCs^19^, backcrossed for >10 generations to C57B6/J using speed congenics, and validated as congenic based on a standardized SNP analysis by Charles River. Considering the 129S ESC origin, the differential expression of genes on chromosome 11 may be a result of the unintended retention of 129S genome proximal to *Nlrp3* during backcrossing.

**Figure 3.**
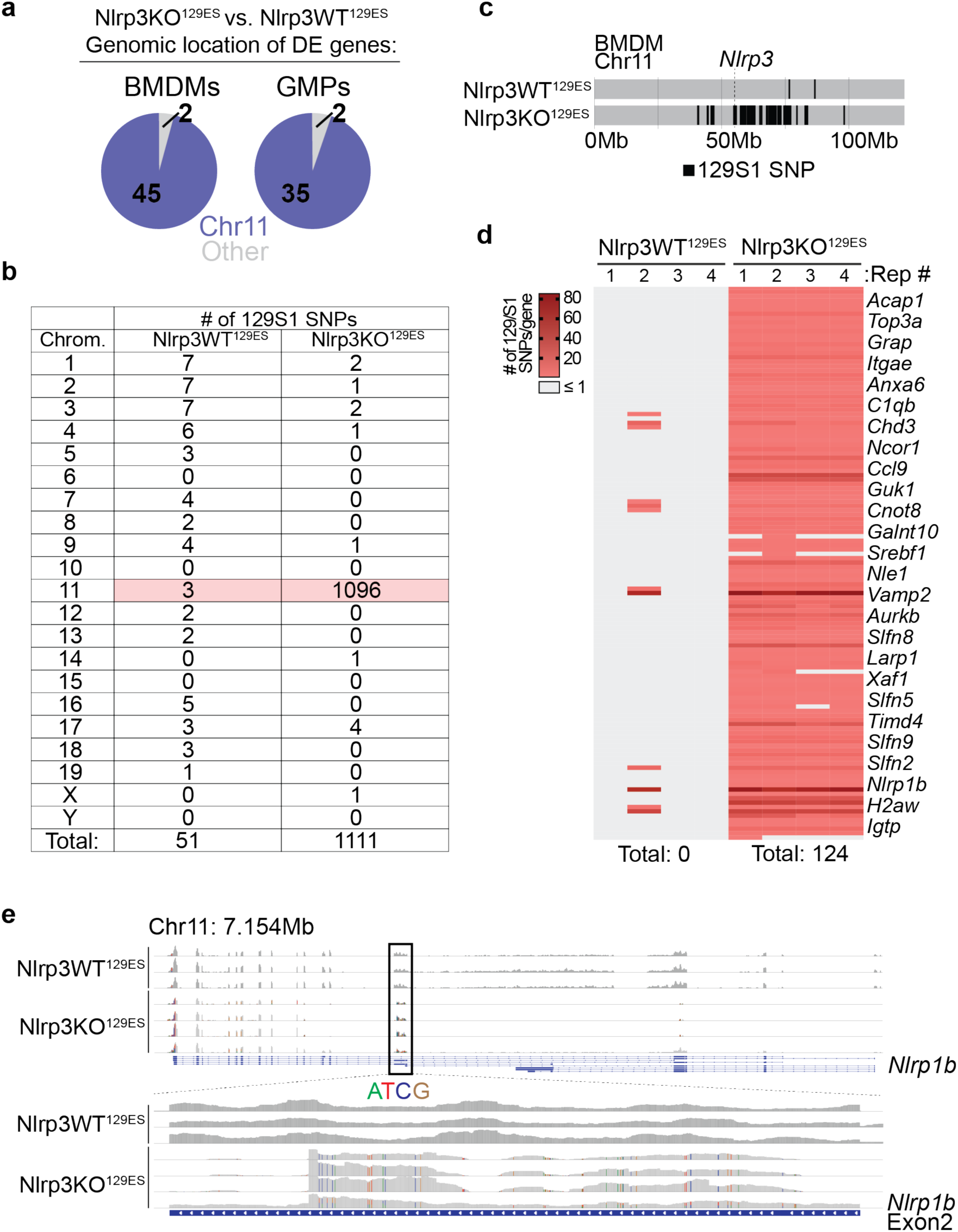
Differentially expressed genes in Nlrp3KO^129ES^ mice are enriched on chromosome 11 and align to the 129S1 mouse strain genome, including *Nlrp1b*. **a.** Pie charts showing the number of differentially expressed (DE) genes (adj. *P* < 0.05) in BMDMs and GMPs between Nlrp3WT^129ES^ and Nlrp3KO^129ES^ mice, and whether they are located on chromosome 11 (blue) or an alternative chromosome (grey). **b.** Table showing the number of identified 129S1 SNPs in at least 3 out of 4 replicates in both Nlrp3WT^129ES^ and Nlrp3KO^129ES^ BMDMs. Values for chromosome 11 are highlighted in red. **c.** Line plot showing every identified SNP mapping to the 129S1 genome compared to the reference genome in RNAseq data from Nlrp3WT^129ES^ and Nlrp3KO^129ES^ BMDMs on chromosome 11. Based on presence in 3 out of 4 biological replicates in each genotype. **d.** Heatmap showing number of 129S1 SNPs in genes located on chromosome 11 from both Nlrp3WT^129ES^ and Nlrp3KO^129ES^. Right: subset of genes identified as containing ≥ 2 129S1 SNPs. Bottom: total number of genes containing ≥ 2 129S1 SNPs in both Nlrp3WT^129ES^ and Nlrp3KO^129ES^ BMDMs. **e.** Snapshot from the Integrative Genomics Viewer (IGV) showing RNAseq reads of *Nlrp1b* from Nlrp3KO^129ES^ and Nlrp3WT^129ES^ mouse BMDMs at baseline. Top: read coverage over the whole *Nlrp1b* gene (expression range: 0 – 300). Bottom: zoom view of reads at exon 2 of *Nlrp1b* (expression range: 0 – 100). SNPs are highlighted in separate colours (adenine (A) = green, thymine (T) = red, cytosine (C) = blue, guanine (G) = brown).

We performed variant calling analysis on polyA RNAseq reads from Nlrp3KO^129ES^ BMDMs and identified 1111 SNPs matching the 129S1 genome in at least 3 out of 4 biological replicates, with 1096 SNPs located on chromosome 11 (**Fig 3B, Supplemental Fig 3A**). By contrast, only 51 such SNPs matching the 129S1 genome were found in RNAseq data from Nlrp3WT^129ES^ BMDMs, with 3 SNPs on chromosome 11 (**Fig 3B, Supplemental Fig 3B**). The vast majority of SNPs identified on chromosome 11 in Nlrp3KO^129ES^ BMDMs were located in close proximity of *Nlrp3* (**Fig 3C**).

As variant calling was performed using RNAseq reads generated from polyA enriched mature RNA, all SNPs identified are located in the coding regions of mRNA from genes expressed in BMDMs. In total, 124 genes located on Chromosome 11 in Nlrp3KO^129ES^ BMDMs contained ≥ 2 SNPs, in at least 3 out of 4 biological replicates (**Fig 3D; Supplemental Table 4**). These genes spanned a region of ∼40Mb, home to a total of 715 genes. No genes matching to 129S1 were identified on chromosome 11 of Nlrp3WT^129ES^ BMDMs using the same criteria (**Fig 3D; Supplemental Table 5**). Genes of 129S origin expressed in Nlrp3KO^129ES^ BMDMs included those previously identified as significantly deregulated, including *Anxa6*, *Xaf1*, and *H2aw* (**Fig 3D**). Furthermore, multiple genes critical for immune cell function were found to contain 129S1 SNPs, including *C1qbp*, *Ccl9*, *Igtp*, *Timd4,* and critically, *Nlrp1b*, defining them as of 129S mouse strain origin (**Fig 3D, E**). Similar results were observed in GMPs, where 956 SNPs matching the 129S1 genome were observed, with ≥ 2 129S1 SNPs within 120 genes (**Supplemental Table 6**).

We extended our analysis to elucidate whether expression changes of genes located on chromosome 11 in Nlrp3KO^129ES^ mice were not just limited to cultured BMDMs and their precursors. Gene expression analysis of microglia purified from adult mouse brains of Nlrp3KO^129ES^ and Nlrp3WT^129ES^ litter-mate controls also revealed a significant deregulation of multiple genes located on Chromosome 11, including *Ubb*, *Ccl4*, *Ccl3*, *Pttg1*, *Xaf1* and again *Nlrp1b* (**Supplemental Fig 4; Supplemental Table 7**).

These data show that Nlrp3KO^129ES^ mice contain a substantial region of 129S mouse strain genome on chromosome 11, in the region surrounding *Nlrp3*, as a result of their production in 129S ESCs, despite extensive backcrossing and validation as congenic. The presence of any identified SNP in protein coding regions may affect the regulation of gene and/or protein expression as well as affect protein function, especially if they are located in functionally critical sites such as at the interfaces of protein-protein interactions.

### NLRP1 hypersensitivity is caused by the 129S *Nlrp1b* allele, and not Nlrp3 deficiency

In order to determine whether the NLRP1 hypersensitivity observed in Nlrp3KO^129ES^ BMDMs was due to NLRP3 deficiency or the presence of the 129S *Nlrp1b* allele, we evaluated NLRP1 inflammasome activation in alternative models of NLRP3 deficiency or loss of function.

MCC950 is a highly selective and potent inhibitor of NLRP3 activation. We treated BMDMs with MCC950 for 7 days in order to mimic long-term loss of NLRP3 function, but did not observe an increase of cell death in response to LPS and Talabostat induced NLRP1 activation (**Fig 4A**). In contrast, cell death in response to NLRP3 activation by LPS and Nigericin was completely abrogated (**Fig 4A**).

**Figure 4.**
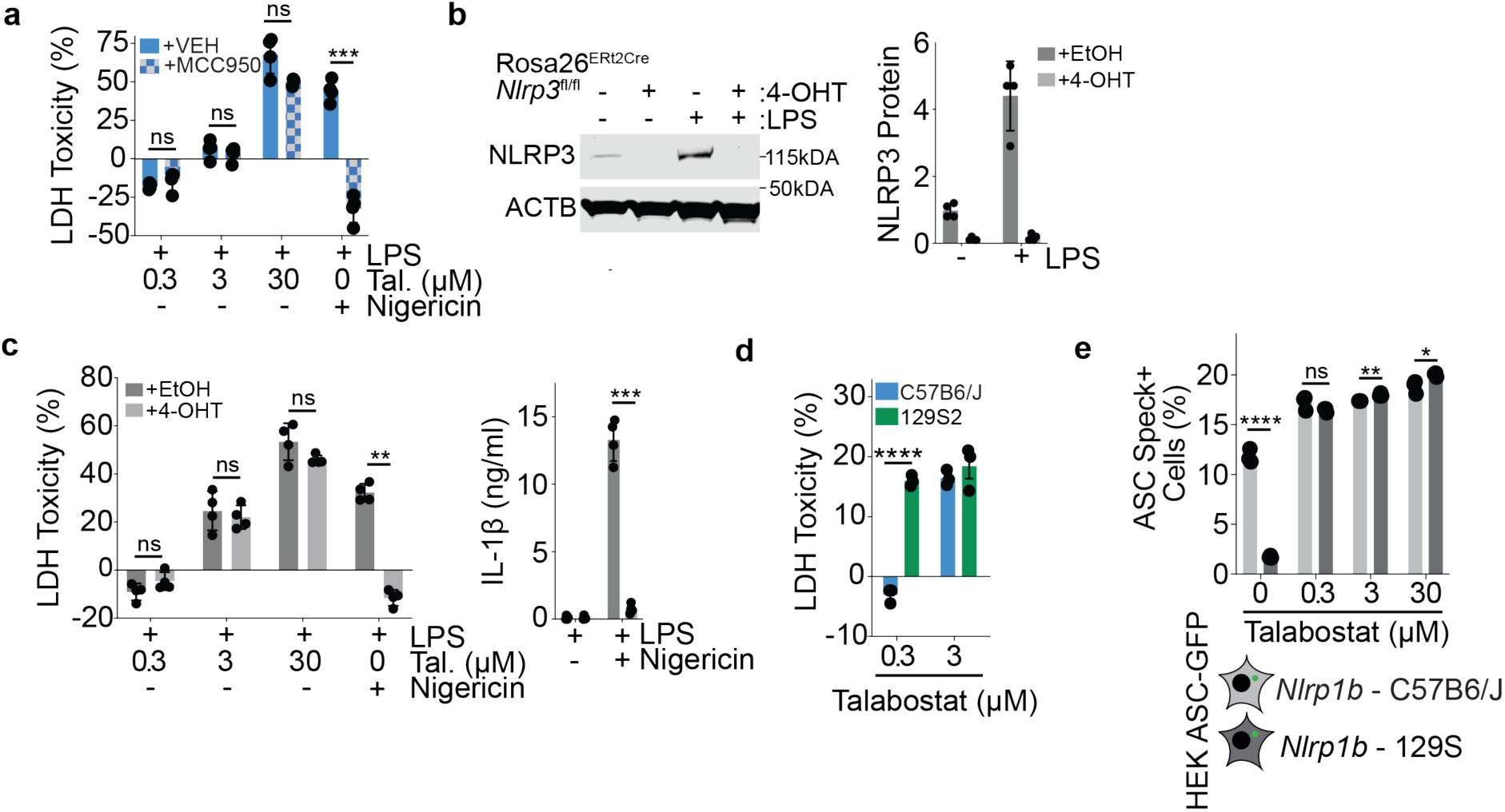
Expression of the 129S mouse strain NLRP1b, not NLRP3 deficiency, drives Nlrp3KO^129ES^ hypersensitivity to Talabostat stimulation. **a.** LDH release, relative to media only total lysis controls, in BMDMs from C57B6/J mice +/−MCC950 after stimulation with Talabostat (0.3 μM, 3 μM or 30 μM) for 24 hours. **b.** Analysis of NLRP3 expression in Rosa26^ERt2Cre^ *Nlrp3*^fl/fl^ BMDMs treated with EtOH or 4-OHT, +/− LPS. Left: Western blot of NLRP3 and ACTB. Right: quantification of NLRP3 protein expression relative to ACTB. **c.** Analysis of LDH and IL-1β release in Rosa26^ERt2Cre^ *Nlrp3*^fl/fl^ BMDMs treated with EtOH or 4-OHT, and co-stimulated with LPS and Talabostat, or LPS + Nigericin. Left: LDH release, relative to total lysis controls. Right: Quantification of IL-1β supernatant concentration. **d.** LDH release, relative to total lysis controls, in BMDMs from C57B6/J (blue) and 129S2 (green) mice after stimulation with Talabostat (0.3 μM or 3 μM) for 24 hours. **e.** Bar plot showing % of ASC speck positive HEK 293 cells stably expressing either 129S or C57B6/J NLRP1b stimulated with Talabostat (0.3 μM, 3 μM or 30 μM) for 16 hours. **a – c**. *P* values were calculated using multiple paired parametric t-tests. FDR (*q*) was calculated using Benjamini Hochberg correction. **d, e**. *P* values were calculated using multiple unpaired parametric t-tests. FDR (*q*) was calculated using Benjamini Hochberg correction. **a – e**. * = *q* < 0.05, ** = *q* < 0.01, *** = *q* < 0.001, **** = *q* < 0.0001, ns = not significant (*q* > 0.05). Error bars represent standard deviation.

While MCC950 treatment prevents NLRP3 activation, as of yet undescribed inflammasome independent NLRP3 functions may not be affected. In order to eliminate NLRP3 expression we utilized a previously unpublished conditional *Nlrp3* allele in combination with tamoxifen inducible ERt2Cre^24^ (Rosa26^Ert2Cre^ *Nlrp3*^fl/fl^). Critically, *Nlrp3*^fl/fl^ mice were generated using C57B6/N ESCs, and as such have a C57B6 *Nlrp1b* allele, unlike the previously described Nlrp3KO^129ES^ mice.

NLRP3 protein expression was significantly reduced in BMDMs following Ert2Cre activation by 4-hydroxytamoxifen (4-OHT) both at steady-state and after LPS treatment (**Fig 4B**). As expected, *Nlrp3* deficient Rosa26^Ert2Cre^ *Nlrp3*^fl/fl^ BMDMs showed a highly significant reduction in LDH and IL-1β release in response to LPS priming and Nigericin stimulation (**Fig 4C**). However, they did not display any differences in LDH release in response to Talabostat (**Fig 4C**). In contrast, BMDMs purified from 129S2 mice, containing the 129S *Nlrp1b* allele, released significantly higher amounts of LDH upon NLRP1 activation, phenocopying the hypersensitivity observed in Nlrp3KO^129ES^ BMDMs (**Fig 4D**).

Furthermore, HEK cells expressing ASC-GFP and equal amounts of either C57B6/J or 129S NLRP1b (Ref. ^25^) showed minor or no differences in ASC speck formation in response to Talabostat (**Fig 4E**). However, as previously reported^25^, C57B6/J NLRP1b expressing HEK cells displayed spontaneous speck formation even in the absence of any triggers.

We conclude that the hypersensitivity observed in Nlrp3KO^129ES^ BMDMs is not due to NLRP3 deficiency or an intrinsic increase in 129S NLRP1b sensitivity, but likely due to increased NLRP1b protein expression in Nlrp3KO^129ES^ BMDMs. Furthermore, strain specific sequence differences may cause C57B6/J NLRP1b to more readily form inflammasomes in the absence of stimulation, requiring its expression to be restricted.

### Nlrp3KO^129ES^ mice misexpress the canonical macrophage marker TIM4

The peritoneal cavity macrophage population can be broadly separated into two functionally distinct subsets: short-lived monocyte-derived macrophages and long-lived tissue resident macrophages. Monocyte-derived macrophages are inflammatory and invade the peritoneum during inflammation, while long-lived tissue resident macrophages regulate tissue homeostasis. In multiple organs including the peritoneal cavity, the cell surface protein TIM4 is used to distinguish long-lived tissue resident macrophages (TIM4+) from short-lived monocyte derived macrophages (TIM4-, **Fig 5A, Supplemental Fig 5,** Ref. ^26–31)^. As such TIM4 is a critical tool in understanding tissue biology and the role of myeloid cells in inflammation.

**Figure 5.**
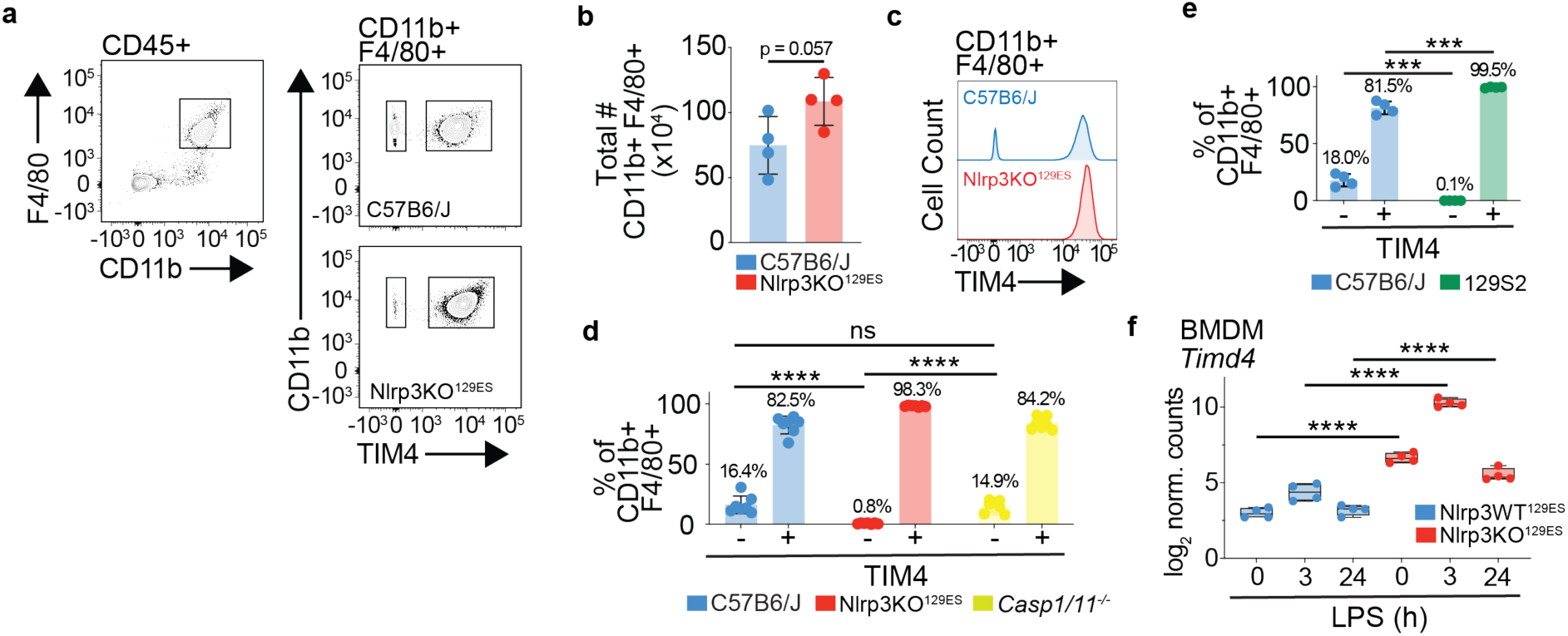
Monocyte derived peritoneal macrophages from Nlrp3KO^129ES^ mice overexpress TIM4. **a.** Representative flow cytometry plots showing identification of TIM4- and TIM4+ macrophages (F4/80+ CD11b+) in C57B6/J and Nlrp3KO^129ES^ mouse peritoneum. Cells were pre-gated on single, live and CD45+ populations. **b.** Bar plot of the total number of macrophages (CD11b+ F4/80+) in each peritoneum of each mouse (Nlrp3KO^129ES^, n = 4; C57B6/J, n = 4). *P* value was calculated using an unpaired t test. Error bars represent standard deviation. **c.** Representative histogram of distribution of TIM4- and TIM4+ macrophages (F4/80+ CD11b+) in C57B6/J and Nlrp3KO^129ES^ mouse peritoneum. **d.** Bar plot of TIM4- and TIM4+ macrophages (F4/80+ CD11b+) in C57B6/J, Nlrp3KO^129ES^ and *Casp1/11*^−/−^ mouse peritoneum. Percentages above bars show the mean value for each group. (Nlrp3KO^129ES^, n = 7; C57B6/J, n = 7; *Casp1/11*^−/−^, n = 8). Adj. *P* values were calculated using a 2-way ANOVA with Turkey’s multiple comparison testing. * = adj. *P* < 0.05, ** = adj. *P* < 0.01, *** = adj. *P* < 0.001, **** = adj. *P* < 0.0001, ns = not significant (adj. *P* > 0.05). Error bars represent standard deviation. **e.** Bar plot of TIM4- and TIM4+ macrophages (F4/80+ CD11b+) in C57B6/J and 129S2 mouse peritoneum. Percentages above bars show the mean value for each group (C57B6/J, n =4; 129S2, n = 4). *P* values were calculated using multiple unpaired parametric t-tests. FDR (*q*) was calculated using Benjamini Hochberg correction. * = *q* < 0.05, ** = *q* < 0.01, *** = *q* < 0.001, **** = *q* < 0.0001, ns = not significant (*q* > 0.05). Error bars represent standard deviation. **f.** log_2_ normalised counts of *Timd4* from RNAseq of Nlrp3KO^129ES^ and Nlrp3WT^129ES^ BMDMs at baseline and after 3h and 24h of LPS (10ng/ml) stimulation (Nlrp3WT^129ES^, n=4; Nlrp3KO^129ES^, n=4). * = adj. *P* < 0.05, ** = adj. *P* < 0.01, *** = adj. *P* < 0.001, **** = adj. *P* < 0.0001, ns = not significant. Wald Test, Benjamini-Hochberg corrected.

Analysis of peritoneal macrophage subtypes revealed that, despite having the same total number of macrophages in the peritoneum (**Fig 5B**), TIM4-macrophages appeared to be virtually absent from Nlrp3KO^129ES^ peritoneal cavity (**Fig 5C**). Quantification of TIM4+ and TIM4-macrophages as a percentage of total macrophages (CD11b+ F4/80+) revealed a significant decrease in the percentage of TIM4-macrophages in the Nlrp3KO^129ES^ peritoneal cavity compared to C57B6/J (16.4% to 0.8%, adj. *P* < 0.0001, **Fig 5D**).

The absence of TIM4-macrophages in Nlrp3KO^129ES^ was not due to the loss of tonic NLRP3 signalling as *Casp1/11^−/−^* mice showed no significant difference in percent frequency of TIM+ and TIM4-macrophages relative to C57B6/J control mice, and significantly more TIM4-macrophages than Nlrp3KO^129ES^ mice as a percent of total macrophages (14.9% to 0.8%, adj. *P* < 0.0001, **Fig 5D**). Strikingly, TIM4-macrophages were also absent from 129S2 mice (**Fig 5E**), suggesting that the lack of TIM4-macrophages in Nlrp3KO^129ES^ mice could be another strain specific effect caused by the retention of 129S ESC derived genome.

RNAseq analysis of Nlrp3ko^129ES^ BMDMs, which are monocyte derived, revealed that the TIM4 coding gene *Timd4* is highly significantly upregulated between Nlrp3KO^129ES^ and Nlrp3WT^129ES^ BMDMs (baseline: log_2_ FC = 7.2, LPS 3h: log_2_ FC = 6.9, LPS 24h: log_2_ FC = 5.0, **Fig 5F**). *Timd4* is located on chromosome 11 (Chr11: 46,808,799) in close proximity to *Nlrp3* (Chr11: 59,539,569). Variant analysis of RNAseq data identified 3 SNPs in *Timd4*, defining it as of 129S origin.

Therefore, monocyte derived macrophages are not absent from the Nlrp3KO^129ES^ peritoneum, but instead aberrantly upregulate the normally tissue resident macrophage restricted marker TIM4 as a result of the presence of 129S genome, confounding cell identification by FACS.

## Discussion

In this study we combine RNAseq and variant calling analysis to identify genetic variation in one of the most frequently used mouse models of NLRP3 deficiency. We found the retention of a ∼40Mb of 129S ESC-derived genomic material proximal to *Nlrp3*, leading to changes in gene regulation, the differential expression of polymorphic proteins, and the NLRP1 inflammasome response.

Post-transcriptional and -translational regulation of NLRP proteins is critical for immune homeostasis, and the upregulation of inflammasome proteins is a key priming step before activation and pyroptosis. We show that the presence of the 129S *Nlrp1b* allele in Nlrp3KO^129ES^ macrophages results in the upregulation of the NLPR1b protein, independently of an increase in *Nlrp1b* gene regulation, sensitizing them to NLRP1 activation by Talabostat. Conversely, Nlrp3KO^129ES^ microglia significantly downregulate *Nlrp1b*. This raises the possibility that microglial phenotypes ascribed to NLRP3 deficiency may be influenced by altered *Nlrp1* expression, especially given that it remains unknown if endogenous murine NLRP1 triggers exist in disease settings such as neurodegeneration.

We further show that C57B6 NLRP1b can form inflammasomes in the absence of any triggers when expressed at the same level as 129S NLRP1b. This suggests that through evolutionary pressure, C57B6 NLRP1b has accrued single nucleotide polymorphisms (SNPs) that restrict its expression in a post-translational manner, protecting mice from spontaneous inflammasome formation. While 129S mice can therefore tolerate higher expression levels of NLRP1b at baseline, this subsequently sensitizes them to NLRP1 inflammasome triggers.

Our findings illustrate a paradigm that can be extended more broadly into the immunology field. X-linked diseases in females, such as Rett Syndrome, display a broad spectrum of severity due to the mosaicism of X inactivation^32,33^. Autoinflammatory skin diseases can arise from heterozygous gain-of-function mutations to NLRP1^34^. Given that immune genes are biased to monoallelic expression^35^, immune related diseases arising from genetic defects may be caused by cells preferentially expressing dominant alleles, undergoing competition that results in the reduction of cells expressing wild-type alleles, or that the expression of a small amount of hyperactive protein is sufficient to drive a phenotype. Heterozygous Nlrp3KO^129ES^ mice, which contain both C57B6 and 129S *Nlrp1* loci, could provide a model for the analysis of allelic bias and dominant inflammatory signalling responses in the context of NLRP1, as well as elucidate novel differences between 129S and C57B6 NLRP1 as described here.

Changes in expression of immune genes are not limited to the *Nlrp1* locus. The upregulation of 129S *Timd4* in Nlrp3KO^129ES^ monocyte-derived macrophages results in their misidentification as tissue-resident macrophages due to increase surface expression of TIM4. As a result, TIM4 cannot be used to determine frequencies of monocyte-derived versus tissue-resident macrophages, and heterogenous cell populations are obtained when purifying cells for molecular analysis such as RNAseq or LC-MS/MS, as well as confounding population analyses by FACS.

Other immune genes identified as containing 129S SNPs include *Igtp*, an interferon induced gene critical for host defense^36^, *C1qbp*, a complement family member that aids the clearance of apoptotic cells^37^, *Itgae*, which codes for the dendritic^38^ and T cell^39^ marker CD103, *Xaf1,* an antagonist of the anti-apoptosis protein XIAP^40^ and the chemokine *Ccl9*, amongst others. Further investigations in Nlrp3KO^129ES^ mice are required to assess the contribution of these differences to immune responses.

Genes required for conserved functions across multiple cell-types were also found to contain 129S SNPs including but not limited to genes related to cell cycle (*Pttg1*, *Aurkb*, *Ccng1*), trafficking (*Gosr1*), histones (*H2aw*), metabolism (*Mat2b, Guk1, Srebf1*, *Galnt10*) translation (*Larp1*, *Mm3*) and transcription (*Cnot8, Top3a*, *Chd3*, *Pol2ra*, *Ncor1*, *Mnt*).

Finally, the region around *Nlrp3* that we detect as being of 129S origin contains genes not expressed in macrophage or GMPs, but of importance in functions such as neuronal signalling (*Gabrg2, Gabra1, Gabra6, Gria1*), olfaction (80 olfactory receptor genes), and development (*Wnt3a*), among others. Further variant calling analysis either on DNA sequencing data or RNAseq from relevant cell-types would be needed to determine the full scale of 129S contamination and the effect on cell function, which could have significant knock-on effects due to alterations in paracrine signalling.

The retention of 129S chromosome 11, including the hypersensitive NLRP1, may affect other *Nlrp3* targeting transgenic mice produced in 129S ESCs. *Nlrp3^A350V^* mice, modelling *NLRP3* human gain-of-function mutations^41^, display skin inflammation in the absence of *Nlrp3^A350V^* expression^41^. Gain-of-function mutations in human *NLRP1* leads to inflammatory skin disease^34^. The unexplained skin inflammation in *Nlrp3^A350V^*mice may therefore be due in part to the presence of 129S-derived hypersensitive NLRP1b.

In transgenic mice, ESC-derived genome retention represents a minority, but potentially functionally important, proportion of the genome. Whole exome sequencing does not inform researchers on which mutated genes are expressed, and to what degree. Our approach to identify genetic heterogeneity in transgenic mice using variant calling analysis on RNAseq data allows for the detection of SNPs in coding regions of functionally relevant expressed genes. Furthermore, it allows for the post hoc analysis of genetic heterogeneity in historical samples that utilized RNAseq. Our identification of 129S1 genome in Nlrp3KO^129ES^ mice, and changes in gene expression and protein function, highlights the careful consideration that should be given to ascribing phenotypes in Nlrp3KO^129ES^ mice to NLRP3 deficiency. Further work should evaluate the full scale of disruption caused by 129S genomic retention, as many other cell-types may be altered by genetic differences or paracrine activities from affected cells. Use of the conditional *Nlrp3* allele we describe here will enable cell-type specific studies of NLRP3 function in homeostasis and disease, independently of confounding SNPs and other polymorphisms.

## Supporting information

ST1. NLRP3ko NLRP3wt BMDM LCMSMS

ST2. BMDM RNAseq

ST3.Nlrp3ko vs. WT GMP RNAseq

ST4. NLRP3ko Macrophage 129S SNPs

ST5. NLRP3WT Macrophage 129S SNPs

ST6. NLRP3ko GMP 129S SNPs

ST7. Microglia RNAseq

## Acknowledgements

We would like to thank Florian I. Schmidt (University Hospital Bonn), Lea Jenster (University Hospital Bonn) and Dagmar Wachten (University Hospital Bonn) for their manuscript proof reading. We would like to thank Dr. Tobias Dierkes (University Hospital Bonn), Dr. Susanne Schmidt (University Hospital Bonn) and the NGS Core Facility of the Medical Faculty at the University of Bonn/WGGC-Bonn for providing support and instrumentation for the 3’mRNA Seq. We would like to thank the Flow Cytometry Core Facility of the Medical Faculty at the University of Bonn for providing support and instrumentation funded by the Deutsche Forschungsgemeinschaft (DFG, German Research Foundation) **–** Project # 216372545.

This work was funded by the European Union’s Horizon 2020 research and innovation program under grant agreement No. 848146 (To_Aition) (to E.L.), by the Deutsche Forschungsgemeinschaft (DFG, German Research Foundation) under Germany’s Excellence Strategy – EXC2151 – 390873048 (to E.L. and F.M.), by the DFG SFB1454 - 432325352 (to E.L. and F.M.), SFB1402 – 414786233 (to E.L.), TRR237 – 369799452 (to E.L.), GRK2168 – 272482170 (to E.L.), the Helmholtz-Gemeinschaft, Zukunftsthema ‘Immunology and Inflammation’ (ZT-0027), and the Faculty of Medicine at the University of Bonn BONFOR 1B Postdoc Fellowship Project ID: O-187.0008.1.

## Author Contributions

F.D.W, E.L and F.M conceived the study. F.D.W and Y.A designed experiments. F.D.W, Y.A, H.E.L and A-K.G performed experiments. F.D.W, A.S, F.S and A.B performed data analysis and visualization. F.D.W, E.L and F.M acquired funding. F.W and F.M wrote the manuscript. All authors reviewed and edited the manuscript.

## Methods

### Mice

Mice were housed in specific-pathogen-free conditions. C57B6/J and 129S2/SvPPasCrl mice were obtained from Charles River Laboratories. Nlrp3KO^1293ES^ mice have been previously described^19^, and were maintained in-house on a C57B6/J background, *Casp1/11^−/−^* mice have been previously described^6^, and were maintained in-house on a C57B6/J background. *Nlrp3*^fl/fl^ (strain ID: 12809) mice were obtained from Taconic. Rosa26ERt2Cre mice have been previously described^24^, and were obtained from Jackson Laboratories (strain ID: 008463). All animal experiments requiring ethical approval were performed under the ethics license AZ. 81-02/04.2019.A336, approved by the ethics committee of North Rhein Westphalia. Male and female mice were used, all experiments were sex matched.

### Cell Culture

For bone marrow derived macrophage (BMDM) production, femurs and tibias were obtained from 8 – 12-week-old mice and flushed with DMEM + 10% FBS through a 70 μm filter. Isolated cells were centrifuged (350g, 5 minutes) and resuspended in DMEM + Glutamax, supplemented with 10% FBS, 1% P/S and 15-20% L929 cell-conditioned medium. BMDMs were then differentiated over a period of 7 days (d) in a cell culture incubator (37°C, 5% CO_2_). Cell culture media was supplemented with an additional 10% of L929 cell-conditioned media on day 3. On the final day of differentiation BMDMs were harvested by cell scraping and resuspended at the desired concentration in DMEM + Glutamax, supplemented with 10% FBS and 1% P/S. For stimulation experiments BMDMs were plated in 96-well plates at a density of 10 x 10^5^ per well.

HEK293 cells stably expressing GFP fused ASC and murine Nlrp1b alleles (a kind gift from Florian I. Schmidt, University Hospital Bonn, DE) were cultured in DMEM + Glutamax, supplemented with 10% FBS and 1% P/S.

### Cell Stimulation

BMDMs were plated at a density of 1 x 10^5^ cells per well of a 96 well plate, or 1 x 10^6^ cells per well of a 6 well plate. For RNAseq and LC-MS/MS experiments, BMDMs were cultured in DMEM + Glutamax supplemented with 1% P/S and 10% FCS, and stimulated with LPS-EB Ultrapure (10 ng/ml, Invivogen) for 0, 3 or 24 hours. For NLRP1 and NLRP3 inflammasome activation, BMDMs were cultured in DMEM + Glutamax supplemented with 1% P/S and 2% FCS. NLRP1 inflammasome activation by Talabostat (Hoelzel) was induced in BMDMs by the simultaneous treatment with LPS (10 ng/ml) and Talabostat (0.3 μM, 3 μM or 30 μM) or a DMSO control, for 16 or 24 hours, either in presence or absence of VX-765 (50 μM), all as indicated. For long-term inhibition of MCC950 treated BMDMs, cells were cultured in DMEM + Glutamax supplemented with 1% P/S and 10% FCS, supplemented every 48 hours with MCC950 (5 μM, Invivogen). NLRP3 inflammasome activation by Nigericin in BMDMs required a priming step where BMDMs were incubated with LPS (10 ng/ml) for 3 hours. Subsequently, BMDMs were further incubated with Nigericin (8 μM, Invivogen) for an additional 90 minutes.

HEK cells were plated at a density of 5 x 10^5^ per well of a 12 well plate. NLRP1 inflammasome formation was activated by incubation with Talabostat (0.3 μM, 3 μM or 30 μM) or a DMSO control for 16 hours in DMEM + Glutamax, supplemented with 10% FBS and 1% P/S.

### Immunoblot

Whole cell extracts were lysed in 1X NuPage LDS Sample Buffer (ThermoFisher), and Protein was loaded in each lane of a 4 – 12% Bolt Bis-Tris Plus gel (ThermoFisher), and was then electrophoretically separated, immunoblotted, and visualised on a LI-COR Odyssey Instrument. Primary antibodies used were NLRP3 (Cryo-2, AdipoGen Life Sciences) and β-Actin (926-42210, LI-COR) were used. Secondary antibodies IRDye 800CW and IRDye 680RD (LI-COR) were used.

### Mouse IL-1β Measurements by HTRF

IL-1β concentrations in cell supernatants were measured by a homogenous time-resolved fluorescence (HTRF) ‘sandwich’ antibody-based assay, following manufacturer’s instructions (62MIL1BPEG, CisBio). Briefly, the anti-mouse IL-1β solutions were mixed at a 1:1 ratio. A portion of 4 μl per well of this mixture was distributed in white low-volume medium-binding HTRF-adapted 384-well assay plates (784075, Greiner Bio-One). This was followed by the addition of the samples (tissue culture supernatants; 16 μl per well). The plates were centrifuged at RT, 1,000 g for 5 minutes, followed by a 3h incubation at RT. HTRF signals were measured using Spectramax i3.

### LDH Cytotoxicity Assay

LDH in cell supernatants was measured using the LDH Cytotoxicity Kit (TaKaRa) according to manufacturer’s instructions. LDH values were normalized to a total lysis control after substraction of spontaneous background signal. Samples were measured on a SpectraMax i3.

### Peritoneal Cavity Cell Isolation for FACS Analysis

Mice were sacrificed by cervical dislocation, and the peritoneal cavity filled with 10ml PBS + 2 mM EDTA. Mice were subsequently shaken to dislodge residing cells, before the PBS + 2mM EDTA was removed. Cells were then centrifuged (350g, 5 minutes) and resuspended in 1X Red Blood Cell Lysis solution (555899, BD Bioscience) for 15 minutes at RT. Cells were then centrifuged again (350g, 5 minutes) and processed for staining.

### Isolation of Bone Marrow Cells for GMP Isolation

In order to isolate GMPs, femurs and tibias were obtained from 8 – 12-week-old mice and flushed with PBS supplemented with 0.5% BSA and 2 mM EDTA (FACS buffer) through a 70 μm filter. Isolated cells were centrifuged (350g, 5 minutes) and resuspended in 1X Red Blood Cell Lysis solution (555899, BD Bioscience) for 15 minutes at RT. Cells were then centrifuged again (350g, 5 minutes) and processed for staining.

### Cell Staining for FACS Sorting and Analysis

All staining was performed at 4°C in PBS + 1% BSA + 2mM EDTA. Bone marrow was stained with the following markers for the isolation of GMPs. Lineage markers, all FITC (B220 (11-0452-85, eBioscience) CD19 (11-0193-85, eBioscience), CD11b (11-0112-82, eBioscience), CD3e (11-0033-82, eBioscience), TER-119 (11-5921-85, eBioscience), CD2 (11-0021-85, eBioscience), CD8b (11-0083-85, eBioscience), CD4 (11-0042-85, eBioscience), Ly-6G (553127, BD Pharmingen)), Sca1-Pacific Blue (108120, BioLegend), c-Kit-APC/Cy7 (47-1172-82, eBioscience), CD16/32-PerCP/Cy5.5 (560540, BD Pharmingen), and CD34-AF647 (128606, BioLegend).

For identification of peritoneal cavity cell subsets the following markers were used: CD45-PE/Cy7 (552848, BD Biosciences), CD11b-BV510 (101245, BioLegend), F4/80-APC (123116, BioLegend), and TIM4-PE (130005, BioLegend). Cells were stained in the presence of Fc Block (553142, BD Biosciences).

For ASC speck staining the following markers were used: ASC-PE (653903, BioLegend). Cells were permeabilized with the FoxP3 Transcription Factor Staining Set (00-5523-00, ThermoFisher) according to manufacturer’s instructions before staining. Cells were stained in the presence of Fc Block (553142, BD Biosciences).

For viability staining in peritoneal cavity populations, cells were incubated with 7-AAD 15 minutes prior to FACS analysis or sorting. For viability staining in BMDMs LIVE/DEAD Fixable Aqua Dead Cell Stain Kit was used (L34957, ThermoFisher). For cell number analysis, Precision Plus Counting Beads (Biolegend) were used according to manufacturer’s instructions.

FACS analysis was performed on a BD Canto, sorting was performed on a BD Aria III.

### Microglia Purification

Microglia were purified from adult brain tissue using the Neural Tissue Dissociation Kit (130-092-628, Miltenyi Biotec), Myelin Removal Beads II (130-096-433, Miltenyi Biotec) and CD11b Microglia Microbeads (130-093-636, Miltenyi Biotec) according to manufacturer’s instructions. Briefly, mice were deeply anesthetized (Ketamine 240 mg/kg and Xylazine 32 mg/kg bodyweight) and the organs were transcardially perfused with 50 ml cold phosphate-buffered saline (PBS, pH 7.4). The brain was removed and the two hemispheres separated, only one hemisphere was used for downstream sample preparation. Brain tissue was subsequently digested, myelin was removed and microglia isolated by bead-based positive selection.

### RNA Extraction and Sequencing

For BMDMs and GMPs RNA was extracted using Picopure RNA Isolation Kit (Thermofisher) according to manufacturer’s instructions. For microglia, RNA was extracted using RNeasy Mini Kit (Qiagen). Residual DNA was removed using RNAse-Free DNase Set (Qiagen). RNA was assessed for quality and quantity (TapeStation, Agilent).

For GMPs and BMDMs, RNA sequencing libraries were prepared using the NEBNext Ultra RNA Library Prep Kit for Illumina following manufacturer’s instructions (NEB, Ipswich, MA, USA). Briefly, mRNAs were first enriched with Oligo(dT) beads. Enriched mRNAs were fragmented for 15 minutes at 94 °C. First strand and second strand cDNAs were subsequently synthesized. cDNA fragments were end repaired and adenylated at 3’ends, and universal adapters were ligated to cDNA fragments, followed by index addition and library enrichment by limited-cycle PCR. Sequencing libraries were validated using NGS Kit on the Agilent 5300 Fragment Analyzer (Agilent Technologies, Palo Alto, CA, USA), and quantified by using Qubit 4.0 Fluorometer (Invitrogen, Carlsbad, CA).

The sequencing libraries were multiplexed and loaded on the flowcell on the Illumina NovaSeq 6000 instrument according to manufacturer’s instructions. The samples were sequenced using a 2×150 Pair-End (PE) configuration v1.5. Image analysis and base calling were conducted by the NovaSeq Control Software v1.7 on the NovaSeq instrument. Raw sequence data (.bcl files) generated from Illumina NovaSeq was converted into fastq files and de-multiplexed using Illumina bcl2fastq program version 2.20. One mismatch was allowed for index sequence identification.

For microglia, 3’mRNA Seq was performed using the QuantSeq 3’mRNA-Seq Library Prep Kit FWD (Lexogen). Final libraries were pooled and sequenced on an Illumina NovaSeq 6000 device with 1×100bp and 10M reads per sample.

### RNAseq Analysis

RNA-seq datasets were processed with nf-core RNA-seq v3.6 ^42^ pipeline using STAR^43^ for alignment and salmon for gene quantification^44^. The library strandedness parameter was set to forward and the reference was set to GRCm38. Statistical analysis was performed in the R environment^45^ with the Bioconductor R-package DESeq2^46,47^. The Benjamini-Hochberg method was used to calculate multiple testing adjusted p-values. For Microglia dataset, only genes with at least 3 read counts in at least 2 samples and at least 5 read counts in total across all samples were considered for analysis. For granulocyte-monocyte progenitor dataset, only genes with at least 20 read counts in at least 3 samples and at least 60 read counts in total across all samples were considered for analysis. For BMDM dataset, only genes with at least 50 read counts (minCount) in at least 3 samples and at least 150 read counts in total across all samples were considered for analysis. Data visualisation, such as volcano plots and heatmaps, were generated upon VST transferred data^48^, using R-packages ggplot2^49^, ComplexHeatmap^50^ and on Graphpad Prism (v10.0.2).

Mutation calling was done on RNA-seq data with nf-core RnaVar v1.0.0 ^42^ pipeline using STAR^43^ for alignment and GATK4^51^ for variant calling. The group-specific mutations were then identified using isec command from bcftools utilities^52^. Unknown group-specific mutations were removed by overlaying the group-specific mutations with known 129S mutations (Accession Nr: GCA_001624185.1). Gene ontology analysis was performed using gProfiler. Statistical domain scope was defined as all genes or proteins expressed in the given dataset. Significance was determined by Bonferroni correction.

### Sample preparation for mass spectrometry

For proteomics analysis without Anl-enrichment, cells were lysed in SDC buffer (1% sodium deoxycholate (SDC), 10 mM tris(2-carboxy(ethyl)phosphine) (TCEP), 40 mM 2-chloroacetamide (CAA), 100 mM Tris-HCl pH 8.5) heated at 95 °C for 10 min and sonicated to shear DNA. Proteins were digested with trypsin and LysC (1:100 enzyme/protein ratio, w/w) at 37 °C, 1000 rpm overnight. Digests were desalted using in-house-made SDB-RPS StageTips.

Desalted peptides were dried in a vacuum concentrator and resolubilized in 0.1% formic acid. Concentrations were determined using a NanoDrop spectrophotometer and normalized between samples for equal peptide injection.

### LC-MS/MS

LC-MS/MS measurements were performed as previously described^53^. Briefly, peptide mixtures were analyzed with an EASY-nLC 1000 coupled to a Orbitrap Exploris 480 (ThermoFisher Scientific). 300ng of peptides were separated on 50 cm in-house-made 75 µm inner diameter columns, packed with 1.9-µm ReproSil C18 beads (Dr. Maisch GmbH) at a flow rate of 300 nl min^−1^ and 60 °C maintained by an in-house-made column oven.

Samples were analyzed without prefractionation in a single shot measurement with a nonlinear 90 minute gradient. Spectra were acquired with data-independent acquisition using full scans with a range of 300–1650 m/z. Data acquisition was controlled by Xcalibur (version 4.4.16.14, Thermo Fisher Scientific).

### LC-MS/MS Analysis

DIA MS raw files were processed by DIA-NN^54^ (version 1.8) with FASTA digest for library-free search and deep learning-based spectra, RTs, and IMs prediction enabled. Precursor FDR was set to 1%, and default parameters were used with the following changes: The precursor range was restricted to 300–1650 m/z, and the fragment ion range to 200 – 1650 m/z. The “--relaxed-prot-inf” option was enabled via the command line. MBR was enabled, neural network classifier was set to “double-pass mode,” and the quantification strategy to “robust LC (high accuracy).” Spectra were matched against the mouse December 2022 UniProt FASTA database. Protein intensities were normalised by the MaxLFQ^55^ algorithm using an in-house script. Bioinformatic analyses were performed with Perseus^56^ (version 1.6.15.0) and R (version 4.1.2). Before statistical analysis, quantified proteins were filtered for at least four valid values in at least one group of replicates. Samples “805_ko_b” and “812_ko_24” were identified as outliers by principal component analysis and subsequently. The remaining missing values were imputed by random draw from a normal distribution with a width of 0.3 and a downshift of 1.8 relatives to the standard deviation of measured values. Statistical tests and parameters used to evaluate annotation enrichment and significant abundance differences of quantified proteins are specified in the figure legends.

**Supplemental Figure 1.**
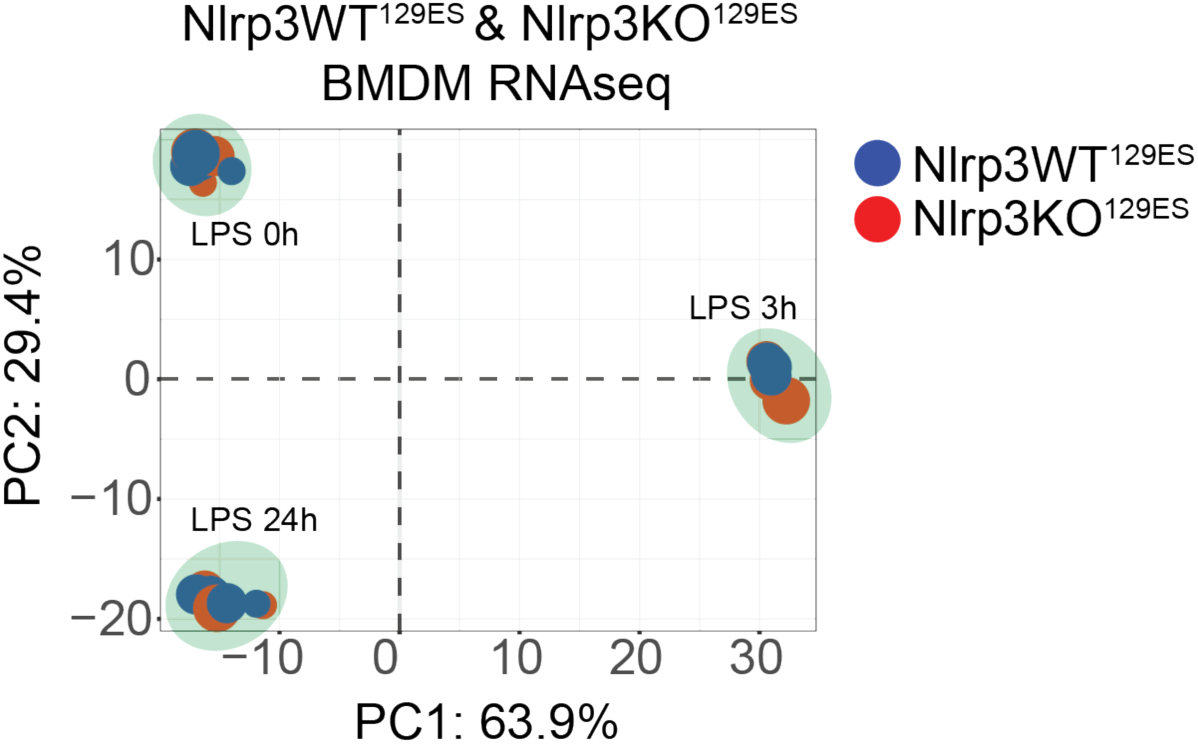
PCA of Nlrp3WT^129ES^ and Nlrp3KO^129ES^ BMDMs RNAseq. Principal component analysis of RNAseq analysis of Nlrp3WT^129ES^ and Nlrp3KO^129ES^ BMDMs at baseline and following LPS stimulation.

**Supplemental Figure 2.**
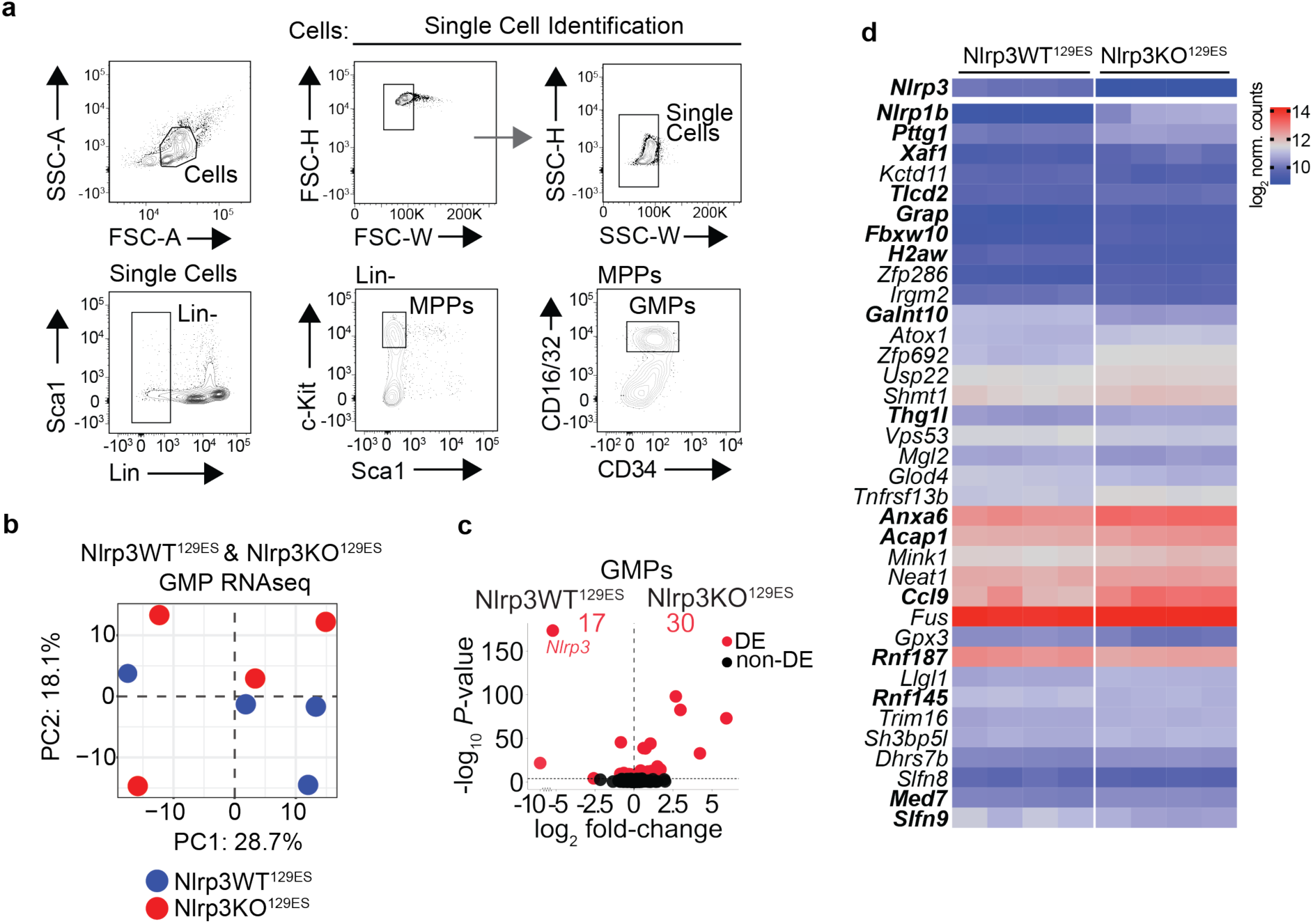
FACS isolation and RNAseq of GMPs from Nlrp3WT^129ES^ and Nlrp3KO^129ES^ mice. **a.** Flow cytometry plots representing gating strategy for sorting of GMPs from mouse bone marrow. **b.** Principal component analysis of RNAseq analysis of Nlrp3WT^129ES^ and Nlrp3KO^129ES^ GMPs. **c.** Volcano plot of gene expression fold-change vs. *P*-value in RNA-seq of Nlrp3WT^129ES^ and Nlrp3KO^129ES^ GMPs. Total number of differentially expressed (DE, adj. *P* < 0.05, Benjamini-Hochberg adjusted) genes, and individual DE genes, are shown in red. **d.** Heatmap of log_2_ normalised counts of significantly differentially expressed (adj. *P* < 0.05) protein coding genes in Nlrp3KO^129ES^ GMPs. Differentially expressed genes shared with Nlrp3KO^129ES^ BMDMs are shown in bold.

**Supplemental Figure 3.**
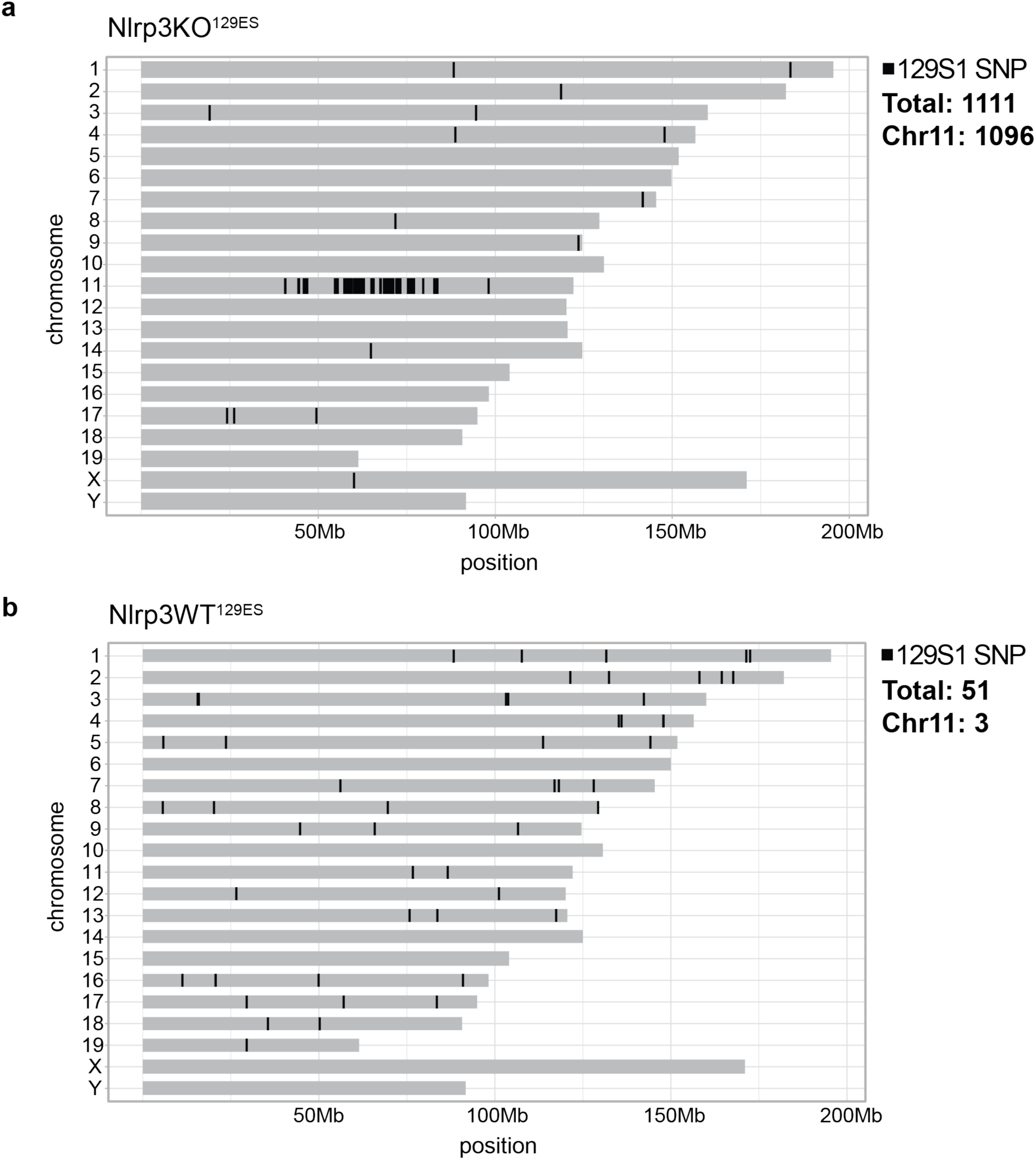
Genomic position of identified 129S1 SNPs in Nlrp3KO^129ES^ and Nlrp3WT^129ES^ BMDMs. **a.** Line plot showing every identified SNP mapping to the 129S1 genome compared to the reference genome in RNAseq data from Nlrp3KO^129ES^ BMDMs on. **b.** Line plot showing every identified SNP mapping to the 129S1 genome compared to the reference genome in RNAseq data from Nlrp3WT^129ES^ BMDMs on. **a-b**. Presence of SNPs based on identification in at least 3 out of 4 biological replicates. Total number of identified SNPs shown on righthand side.

**Supplemental Figure 4.**
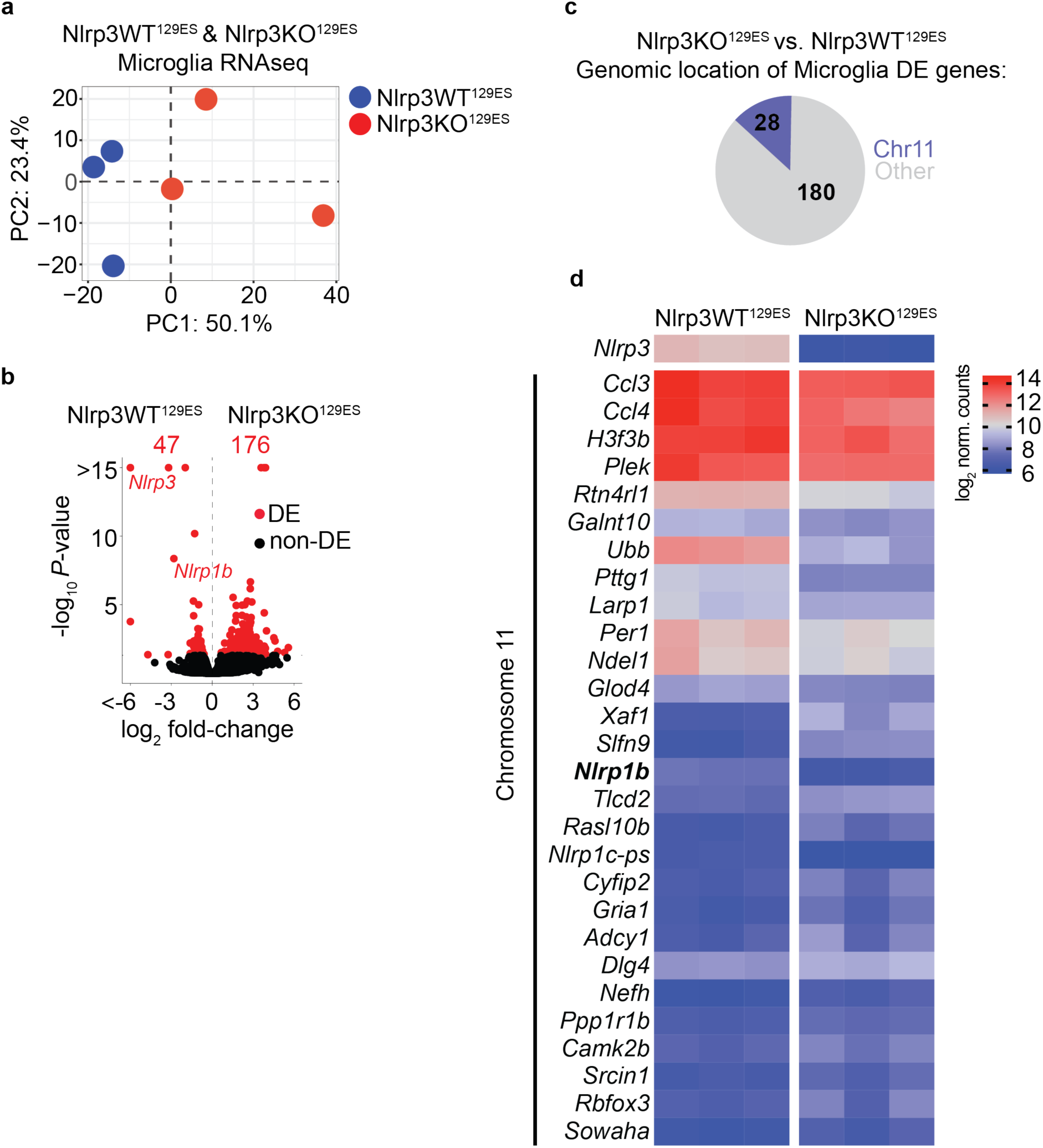
Gene expression analysis of Nlrp3KO^129ES^ and Nlrp3WT^129ES^ microglia. **a.** Principal component analysis of RNAseq analysis of Nlrp3WT^129ES^ and Nlrp3KO^129ES^ microglia. **b.** Volcano plot of gene expression fold-change vs. *P*-value in RNA-seq of Nlrp3WT^129ES^ and Nlrp3KO^129ES^ microglia. Total number of differentially expressed (DE, adj. *P* < 0.05, Wald Test, Benjamini-Hochberg adjusted) genes, and individual DE genes, are shown in red. **c.** Pie chart showing the number of differentially expressed (DE) genes (adj. *P* < 0.05) in microglia between Nlrp3WT^129ES^ and Nlrp3KO^129ES^ mice, and whether they are located on chromosome 11 (blue) or an alternative chromosome (greu). **d.** Heatmap of log_2_ normalised counts of significantly differentially expressed genes (adj. *P* < 0.05) located on Chromosome 11 between Nlrp3WT^129ES^ and Nlrp3KO^129ES^ microglia.

**Supplemental Figure 5.**
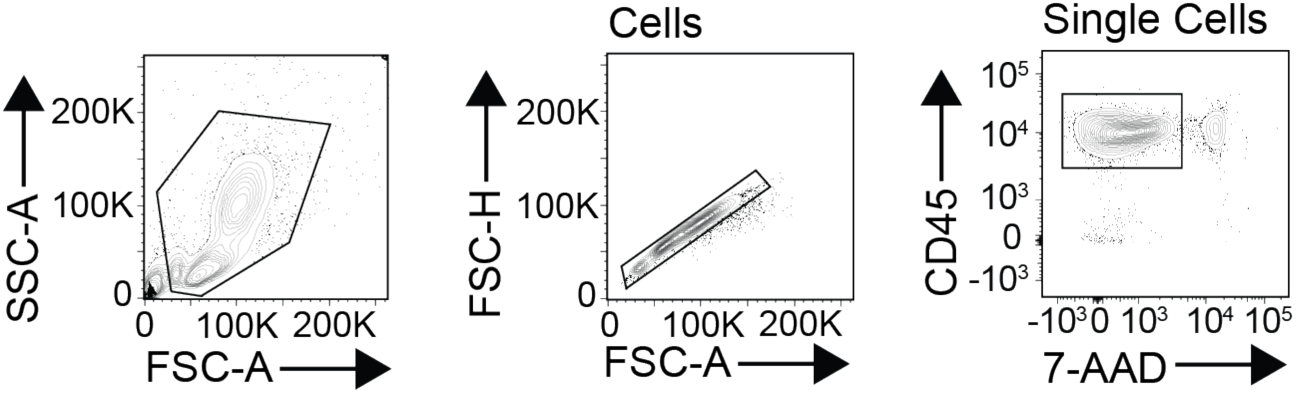
FACS identification of live CD45+ cells in the peritoneum. Flow cytometry plots representing gating strategy for identification of CD45+ Live cells from peritoneal cavity

